# Comprehensive visualization of cell-cell interactions in single-cell and spatial transcriptomics with NICHES

**DOI:** 10.1101/2022.01.23.477401

**Authors:** Micha Sam Brickman Raredon, Junchen Yang, Neeharika Kothapalli, Wesley Lewis, Naftali Kaminski, Laura E. Niklason, Yuval Kluger

**Author notes:** These authors contributed equally (Micha Sam Brickman Raredon, Junchen Yang).

## Abstract

**Summary:** Recent years have seen the release of several toolsets that reveal cell-cell interactions from single-cell data. However, all existing approaches leverage mean celltype gene expression values, and do not preserve the single-cell fidelity of the original data. Here, we present **NICHES** (**N**iche **I**nteractions and **C**ommunication **H**eterogeneity in **E**xtracellular **S**ignaling), a tool to explore extracellular signaling at the truly single-cell level. NICHES allows embedding of ligand-receptor signal proxies to visualize heterogeneous signaling archetypes within cell clusters, between cell clusters, and across experimental conditions. When applied to spatial transcriptomic data, NICHES can be used to reflect local cellular microenvironment. NICHES can operate with any list of ligand-receptor signaling mechanisms and is compatible with existing single-cell packages and pseudotime techniques. NICHES is also a user friendly and extensible program, allowing rapid analysis of cell-cell signaling at single-cell resolution.

**Availability and implementation:** NICHES is an open-source software implemented in R under academic free license v3.0 and it is available at github.com/msraredon/NICHES. Use-case vignettes are available at https://msraredon.github.io/NICHES/.

## 1. Background

Cellular phenotype across tissues and organs is heavily influenced by the biological microenvironment in which a given cell resides (Baccin, et al., 2020; Davidson, et al., 2020; McCarthy, et al., 2020; Nabhan, et al., 2018; Qadir, et al., 2020; Rodda, et al., 2018; Tikhonova, et al., 2020; Zhou, et al., 2018). Understanding the influence of cell-cell signaling on cell phenotype is a major goal in developmental and tissue biology and has profound implications for our ability to engineer tissues and next-generation cellular therapeutics. Single-cell technologies, which capture information both from individual cells and their surrounding cellular environment at the same time, are uniquely suited to exploring phenotype-environment relations. Many techniques are available to extract and prioritize extracellular signaling patterns from single-cell data, reviewed well in (Dimitrov, et al., 2022) and (Armingol, et al., 2021), including CellPhoneDB (Efremova, et al., 2020), NicheNet (Browaeys, et al., 2019), CellChat (Jin, et al., 2021), Connectome (Raredon, et al., 2022), SingleCellSignalR (Cabello-Aguilar, et al., 2020), iTALK (Wang, et al., 2019), iCELLNET (Noël, et al., 2021), Cellinker (Zhang, et al., 2021), CellCall (Zhang, et al., 2021), and PyMINEr (Tyler, et al., 2019). All of these techniques, however, rely on mean expression values calculated from single-cell clusters. Mean expression representation does not take full advantage of the single-cell resolution of the original measurements, thereby obscuring the rich repertoire of signaling patterns between cells. The field can benefit from a tool to assesses cell-cell signaling at the *truly single-cell level*, so that intra- and inter-cluster signaling patterns can be explored within observed data.

Here, we describe NICHES (**N**iche **I**nteractions and **C**ommunication **H**eterogeneity in **E**xtracellular **S**ignaling), a software package to characterize cellular interactions in ligand-receptor signal-space at the single-cell level and to allow cross-platform low-dimensional embeddings of the resulting information. NICHES is designed for analysis of two types of cellular interactions: cell-cell signaling (defined as the signals passed between cells, determined by the ligand expression of the sending cell and the receptor expression of the receiving cell) and cellular niche (defined as the signaling input to a cell, determined by the ligand expression of surrounding or associated cells and the receptor expression of the receiving cell). The outputs from NICHES may be analyzed using existing single-cell software including Seurat (Butler, et al., 2018), Scanpy (Wolf, et al., 2018), Scater (McCarthy, et al., 2017), and Monocle3 (Cao, et al., 2019), thereby allowing deep computational analysis of cell signaling systems topology unapproachable with existing tools.

## 2. Approach

NICHES takes single-cell data as input and constructs matrices where the rows are extracellular ligand-receptor signaling mechanisms and the columns are cell-cell extracellular signaling interactions (Fig. 1A-C). Cell-cell interactions are represented as columns whose entries are created by crossing ligand expression on the sending cell with receptor expression on the receiving cell, for each mechanism (Fig. 1B, see Methods). Cellular niches, an estimate of cellular microenvironment, are represented as columns that are created by crossing mean ligand expression from sets of sending cells with the receptor expression on receiving cells (Fig. 1C, see Methods). Row names are defined by the ground-truth ligand-receptor mechanism list set by the user. NICHES provides built-in access to ligand-receptor lists from the OmniPath and FANTOM5 databases (Ramilowski, et al., 2015; Türei, et al., 2021) and is compatible with custom mechanism lists containing any number of ligand or receptor subunits (Fig. S1, see Methods).

**Figure 1:**
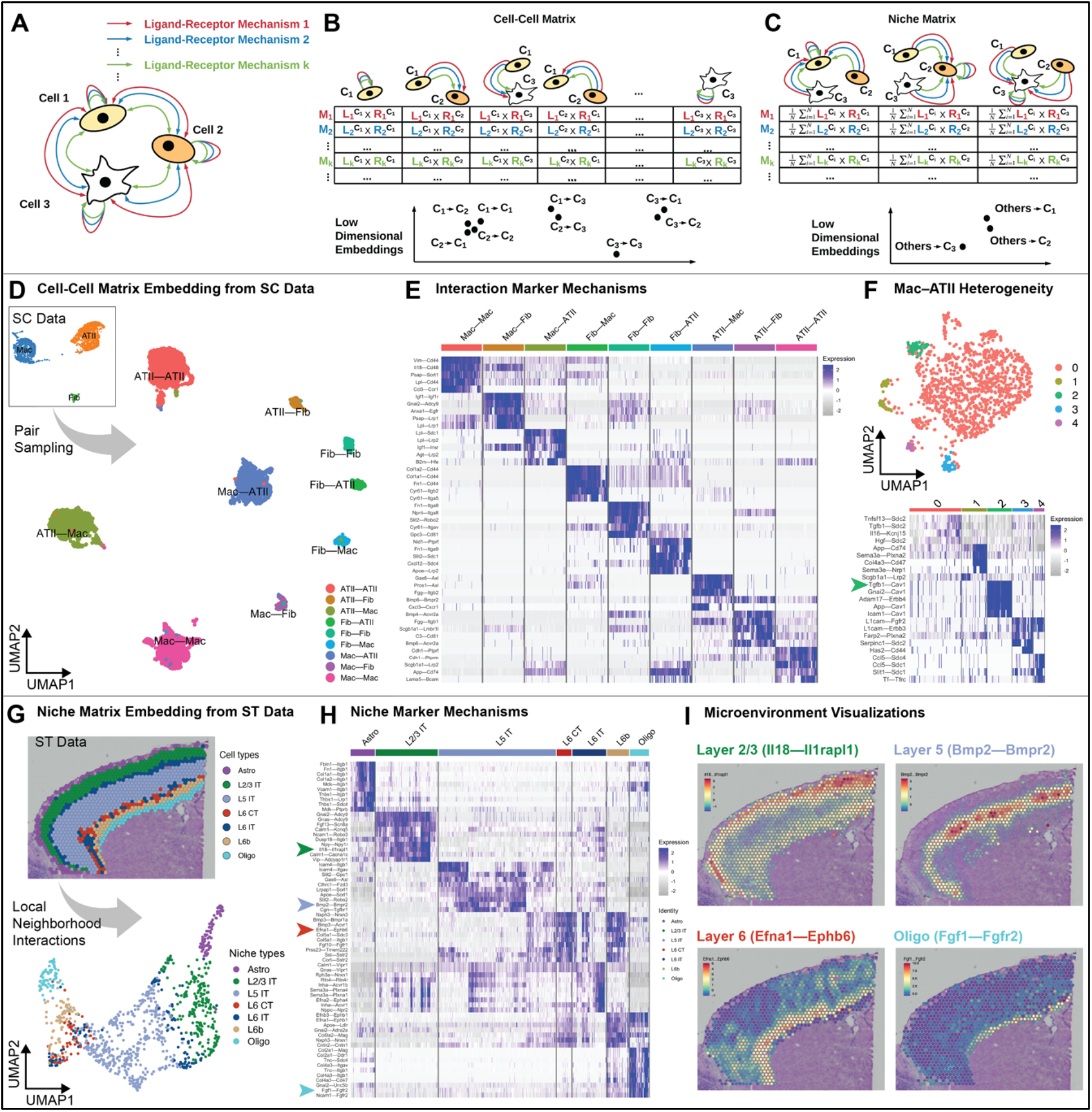
NICHES allows analysis of cell-cell interactions with single-cell resolution. A) A set of cells may interact through many different ligand-receptor mechanisms. B) NICHES represents cell-cell interactions as columns whose entries are calculated by crossing ligand expression on the sending cell with receptor expression on the receiving cell, for each signaling mechanism. Low-dimensional embeddings may then be made of cell-cell interactions. Note schematic clustering of similar profiles. C) Cellular microenvironments, or niches, of each cell are represented as columns calculated by crossing mean ligand expression in the system with receptor expression on the receiving cell. This allows low-dimensional embedding of a proxy for sensed microenvironment for each cell. D) NICHES analysis of single-cell (SC) data of three cell types co-localized in the rat pulmonary alveolus yields a quantitative cell-cell signaling atlas visualized by low-dimensional embedding. E) Biologically-relevant marker mechanisms may be identified for each celltype-celltype interaction. Because single-cell fidelity is preserved, NICHES allows observation of fine intra-cluster heterogeneity unobservable using mean-wise techniques. F) Further analysis of a single celltype-celltype cross allows identification of mechanisms marking only subsets of cell pairings (see Tgfb1-Cav1 in this instance, green arrow, which marks Cluster 2.) G) Local microenvironment may be estimated from spatial transcriptomic (ST) datasets by limiting cell-cell interactions to those within local neighborhoods, yielding a ‘niche’ atlas for each transcriptomic spot, which may be visualized in low dimensional space. H) Signaling mechanisms marking the microenvironments of selected celltypes. Fgf1-Fgfr2 (cyan arrow) is a known potent regulator of oligodendrocyte phenotype (Furusho, et al., 2020; Furusho, et al., 2015) and here is found to be associated with oligodendrocyte-labeled spots. I) Microenvironment mechanisms may be directly visualized in situ.

When applied to spatial transcriptomic data, interactions may be constrained to those occurring between spatial neighbors. When applied to single-cell data, NICHES assumes full cellular connectivity for niche interactions and samples unique cell pairs from each celltype-celltype cross for cell-cell interactions. Biological assumptions, limitations to the mathematical formalism, best practice recommendations, and detailed methods are provided in the Supplement. Replicable vignettes covering a wide-variety of biological use-cases are available both within the NICHES software package and at https://msraredon.github.io/NICHES/.

## 3. Application

### 3.1 Advantages of NICHES over Existing Techniques

Because NICHES does not leverage cluster-wise mean values, signaling heterogeneity hidden by existing cell-cell signaling inference tools is easily observed (Fig. S2A-C, Supplemental Text). Archetype shifts between conditions with conserved mean expression may also be observed, which are difficult to capture with existing tools (Fig. S3A-D, Supplemental Text). NICHES also uniquely allows users to explore changes in system-level signaling due to the addition or loss of cell populations, a task which is not possible with current methods (Fig. S4A-C, Supplemental Text).

### 3.2 Cell-Cell Signaling Atlases

NICHES allows comprehensive visualization of ligand-receptor patterns that are present in single-cell systems data (Fig. 1D-F). A uniform sample is taken of every celltype-celltype interaction resulting in a cell-cell signaling atlas that can be viewed via low-dimensional embedding (Fig. 1D). Celltype-celltype interactions generally display quantifiable signaling signatures as well as intra-relationship heterogeneity (Fig. 1E). Individual celltype-celltype crosses may be subclustered to further explore relationship heterogeneity and to identify mechanisms marking subtypes of cell-cell crosses (Fig. 1F).

### 3.3 Mapping Local Microenvironment in Spatial Atlases

NICHES can estimate local microenvironment in spatial transcriptomic data. Interactions may be limited to spatial nearest neighbors, allowing an estimation of local niche for each transcriptomic spot (Fig. 1G). Celltypes generally display stereotyped niche signatures with observable intra-niche heterogeneity (Fig. 1H). NICHES can reveal tightly localized microenvironments and mechanisms which can be visualized in spatial context (Fig. 1I). Sub-clustering can reveal microenvironment heterogeneity associated with tissue boundaries and transition regions (Fig. S5).

### 3.4 Differential Analysis Across Conditions and Pseudotemporal Orderings

NICHES allows differential and Pseudotemporal analysis of cell-to-cell signaling, system-to-cell signaling, and cell-to-system signaling in both spatial and single-cell datasets. For brevity, we have compiled a series of vignettes online (https://msraredon.github.io/NICHES/) demonstrating these specific use cases. Best practices are discussed in the Supplement.

## 4. Conclusion

NICHES is a simple but powerful approach to explore cell-cell signaling interactions in single-cell and spatial transcriptomic data. NICHES supplements the capabilities of current techniques, allowing single-cell resolution of niche signaling and cell-cell interactions, and establishes rich representations to analyze environment-phenotype relationships in tissues.

## Acknowledgements

M.S.B.R. acknowledges support by NIH grant F30HL143906 and start-up funds from the Yale School of Medicine and the Yale Department of Anesthesiology. Y.K. acknowledges support by NIH grants R01GM131642, UM1DA051410, U54AG076043, P50CA121974, and U01DA053628. This work was supported in part by grant T32GM086287 from the National Institute of General Medical Sciences (NIGMS). The opinions are solely the authors’ and do not necessarily represent the thoughts or opinions of NIGMS, NIH, or the United States government.

## Supplementary Materials for

### Supplemental Text

#### Assumptions, Limitations, and Best Practices

1. NICHES is an extracellular connectivity tool designed to quantify the likelihood of cell-cell communication based on cognate ligand-receptor expression between cells. Complete biological communication requires not only this cognate ligand-receptor expression but also appropriate cellular co-localization for a given mechanism, evidence of signal transduction, absence of downstream signal inhibition, and subsequent phenotypic response, which is cell-state dependent. We think of NICHES as quantifying the mechanistic ‘wavelengths’ that a sending and receiving cell are jointly ‘tuned’ to.
2. In its present form, NICHES multiplies ligand expression on sending cells with receptor expression on receiving cells. This operator was chosen because it is zero-preserving.
3. When mechanisms are queried which containing more than one ligand subunit or more than one receptor subunit, NICHES multiplies the expression of all ligand subunits to yield a single ligand-mechanism expression value and multiplies the expression of all receptor subunits to yield a single receptor-mechanism expression value (see Figure S1 for a diagram of this computation). We have chosen this approach so that zero expression of any subunit within a given mechanism yields zero connectivity for that mechanism. It should be noted, however, that the sparse nature of single-cell transcriptomic data means that NICHES may output zero connectivity for a given mechanism between two cells if all subunits were not captured in a cell used for a given cell-cell pairing. Imputation prior to running NICHES can lessen this artifact, at the cost of using pseudo-values to calculate connectivity. For a demonstration of this difference, please see https://msraredon.github.io/NICHES/articles/02%20NICHES%20Single.html
4. The SystemToCell and CellToSystem outputs estimate connectivity between an entire single-cell system and each individual cell within that system. These tools are designed to quantify changes in system signaling character due either to altered cellular gene expression or to altered cellular distribution or representation. These tools use the mean operator to combine ligand information from multiple sending cells and/or to combine receptor information from multiple receiving cells.
5. When spatial information is provided, NICHES allows users to limit analysis to connectivity within either a defined radius or to local nearest-neighbor communities. However, when spatial information is not provided, NICHES treats all barcodes within an input system as available for sampling and analysis. If cells have been captured which are not to be treated as within a biological system of interest, we recommend removing these cells from the input data prior to running the SystemToCell or CellToSystem functionality, as otherwise ligand or receptor information from these cells will be included downstream.
6. We recommend standardizing total input cell number to maximize the accuracy of cross-system system-level comparisons. We make this recommendation because artifacts may otherwise theoretically be created in instances in which variable cell-number denominators act on constant system-expression numerators for a given mechanism.
7. Batch effects which cause biologically artifactual variance in ligand and receptor expression will affect the outputs from NICHES. This should be noted by users.
8. Differential testing may be performed on NICHES output data. If comparing experimental conditions containing multiple batches, users may run NICHES either for each batch independently or for an entire condition (containing multiple batches) at once. Running NICHES on individual batches will preserve batch effects for downstream analysis. Conversely, running NICHES on an experimental condition containing multiple batches will have the effect of smoothing cell-signaling batch effects, since cells will be crossed with other cells within their condition but not necessarily within their specific batch.
9. If a single dataset is input into NICHES which contains information across multiple experimental conditions, cells may be crossed with cells outside of their respective condition. We currently do not recommend doing this unless such patterns are deliberately of interest to a user.
10. Pseudotime ordering may be performed on either the NICHE output data or on the original cell data and then used to align NICHES information. We currently recommend the latter approach, as we have not investigated how well the NICHES outputs adhere to the fundamental assumptions underlying the diverse array of existing pseudotemporal techniques.

### Supplemental Findings

#### Simulation 1: NICHES reveals hidden intra-cluster signaling heterogeneity

To demonstrate NICHES’s ability to discover intra-cluster heterogeneity, we generated a synthetic single-cell RNA-seq dataset with 400 cells equally divided into 2 cell types (C1 and C2), which can be separated by their marker genes (Figure S2, A, top).

Additionally, there are 2 subtypes S1 and S2 within C1 and 2 subtypes S3 and S4 within C2 (Figure S2, A, bottom). By design, S1 and S2 interact differentially with S3 and S4 through 2 ligand-receptor mechanisms. Specifically, S2 has a higher expression of *A2m* ligand gene compared to S1 (corresponding receptor gene is *Lrp1* that is highly expressed in S3 and S4), and S3 has a higher expression of receptor gene *Agtr2* compared to S4 (corresponding ligand gene is *Ace* which is highly expressed in S1 and S2), as shown in Figure S2, B. Due to the subtle gene expression difference, one cannot distinguish the subtypes based on the overall gene expression profiles. But because the nuance involves ligand and receptor genes, we may apply NICHES to capture this signal.

Figure S2, C, shows the cell-cell matrix output of NICHES, embedded in 2D mechanism space, clustered via k-means (top) and labeled by ground-truth subtype interaction (bottom). Sending cells are all from C1 and receiving cells are all from C2, and the space is 2-dimensional because there are only 2 ligand-receptor mechanisms in this simulation: *A2m-Lrp1* and *Agtr2-Ace*. Note that the clustering results (Figure S2, C, top) are almost completely identical to the ground truth subtype interaction pair labels (Figure S2, C, bottom). For reference, we have added a single black point representing the single mean connectivity value between C1 and C2 which results from using a mean-wise computational method to assess ligand-receptor connectivity in this simulation.

#### Simulation 2: NICHES preserves information regarding differential signaling distributions in disparate experimental conditions

Next, we sought to demonstrate that NICHES can register how cellular archetype shift influences cell-cell connectivity. We simulated 2 scRNAseq cases, both of which contain 2 cell types (C1 and C2), as shown in Figure S3, A. In both cases, we simulated 1 active ligand-receptor channel between C1 and C2 (C1 expresses the ligand gene A2m and C2 expresses the receptor gene Lrp1). C1 expression level of ligand A2m is sparse but high in Case 1, and broad but low in Case 2, with similar mean values (Figure S3, B). By design, the expression level of receptor by population C2 is identical in both cases. Because the mean expression level of A2m is similar, the mean connectivity of A2m-Lrp1 is similar between the 2 cases (Figure S3, C). NICHES, however, is able to detect this significant difference in connectivity (Figure S3, D).

#### Simulation 3: NICHES allows high-dimensional visualization of altered systems-cell topology due to addition or removal of cells

Next, we sought to evaluate the ability of NICHES to map changes in system-to-cell signaling topology. We simulated two cell systems to be compared. The first contains two cell types (C1 and C2, Figure S4, A) while the second contains 3 cell types (C1, C2, and C3, Figure S4, B). C1 and C2 are identical in character in each case, and are connected via two simulated mechanisms, A2m-Lrp1 and Ace-Agtr2, where C1 cells express the ligands A2m and Ace and C2 cells express the receptors Lrp1 and Agtr2. C3, which is only present in the three-cell system, also expresses ligands A2m and Ace. From the perspective of a receiving cell (from population C2, in this simulation), the mean expression of ligand is higher in the three-cell system as compared to the two-cell system. NICHES quantifies this difference and allows analysis and visualization between conditions (Figure S4, C).

### Supplemental Figures

**Figure S1:**
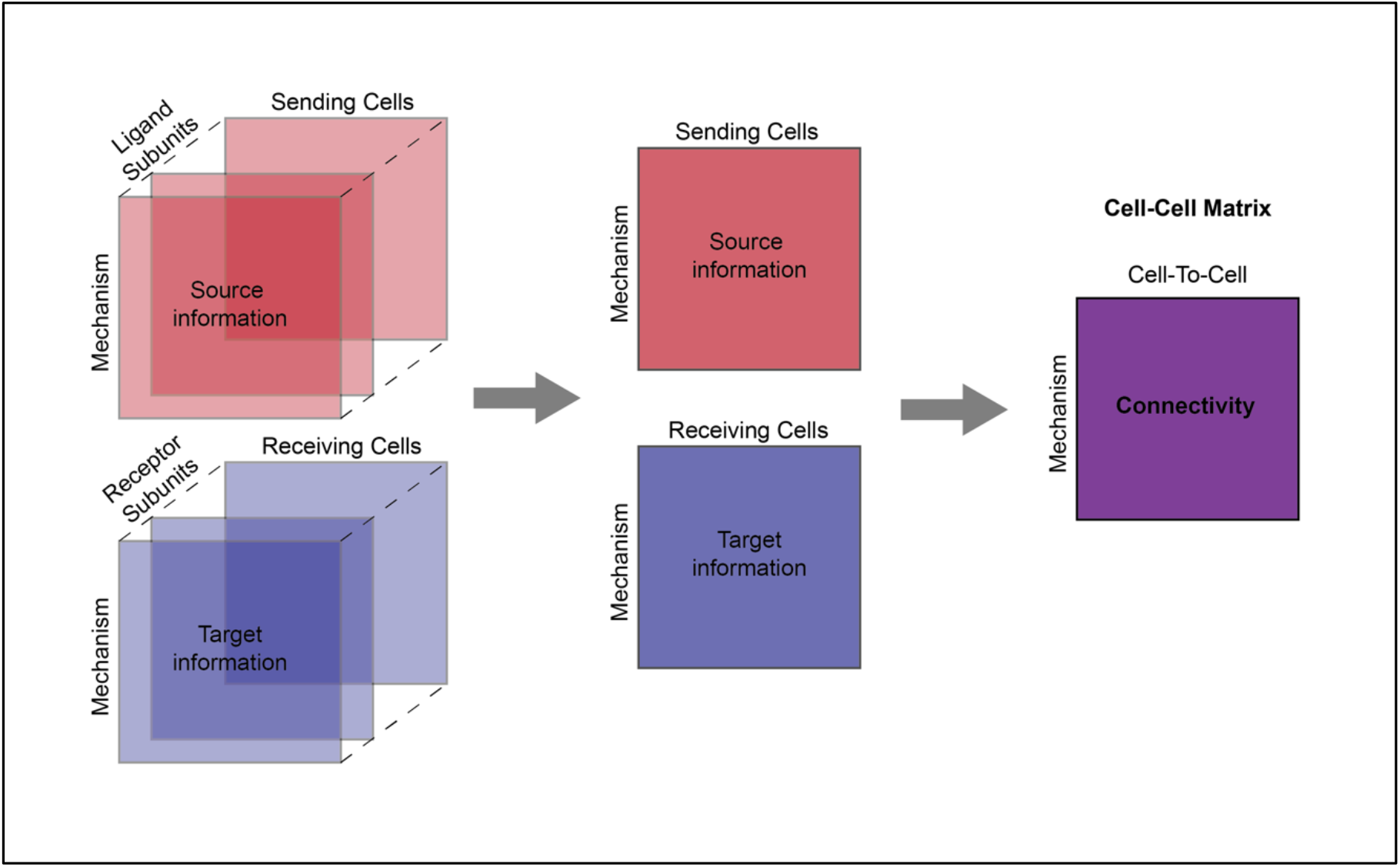
NICHES connectivity for mechanisms with multiple subunits. NICHES is capable of calculating intercellular connectivity for ligand-receptor mechanisms with any number of subunits. For a given mechanism, ligand subunit expressivities on the sending cell are multiplied together and receptor subunit expressivities on the receiving cell are multiplied together. These two values are then multiplied to yield mechanism connectivity between the sending and receiving cell. This operation is zero-preserving by design, so that the lack of expression of even a single subunit on either the sending or receiving cell within a given cell-cell pairing will cause a connectivity value of zero.

**Figure S2:**
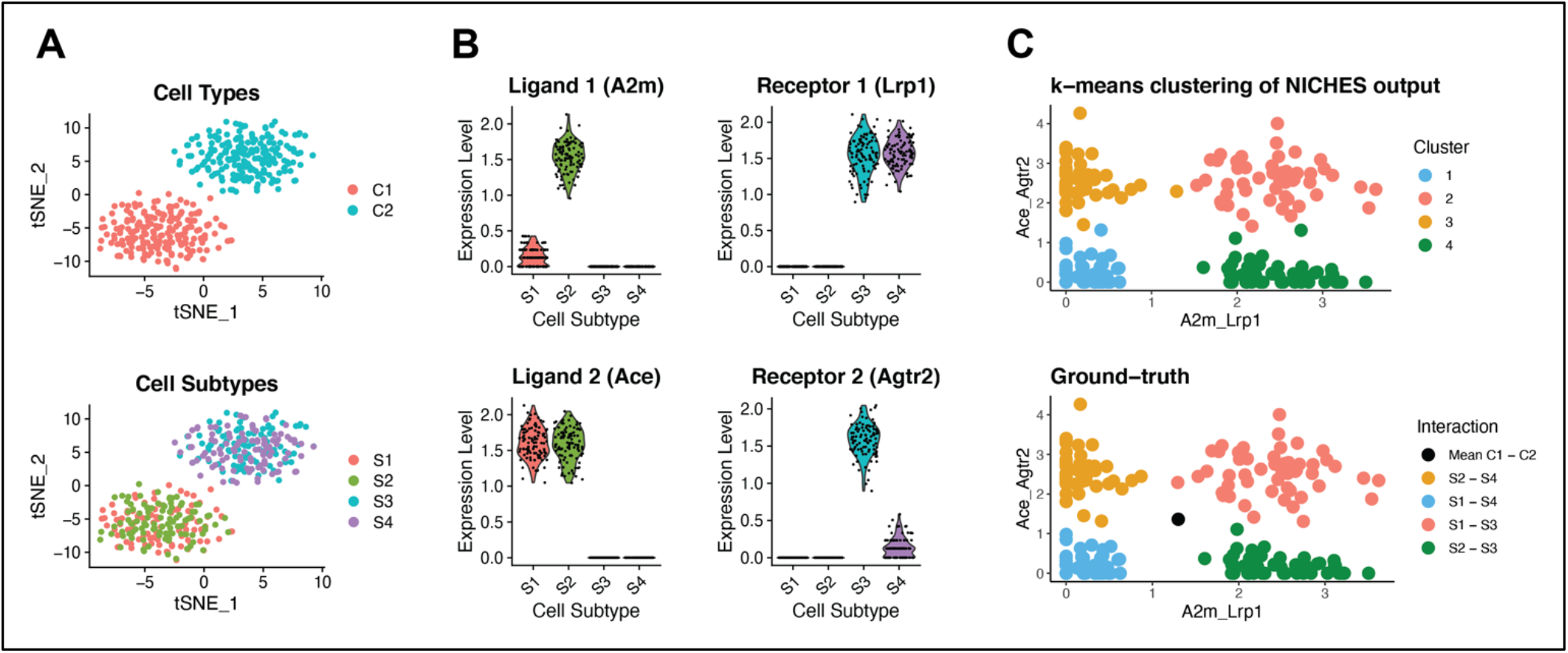
NICHES captures heterogeneity in cell-cell connectivity. In this simulation using synthetic data (see Methods) there are two celltypes labeled C1 and C2 containing signaling subtypes S1-S4 which do not resolve in gene space (A). These subtypes communicate in distinct ways via two distinct signaling mechanisms: A2m-Lrp1 and Ace-Agtr2 (B). Subpopulation S2 expresses ligand A2m higher than S1 while subpopulation S3 expresses receptor Agtr2 higher than S4. This expression pattern creates four distinct cell-cell signaling relationships even though only two celltypes have been crossed. NICHES allows rapid observation of these distinct relationships using two-dimensional embeddings and k-means clustering (C, top) which closely matches the ground truth subtype crosses in this simulation (C, bottom). Mean connectivity between C1 and C2 is represented in black in the lower panel of (C). A biological counterpart to this simulation is show in Figure 1F.

**Figure S3:**
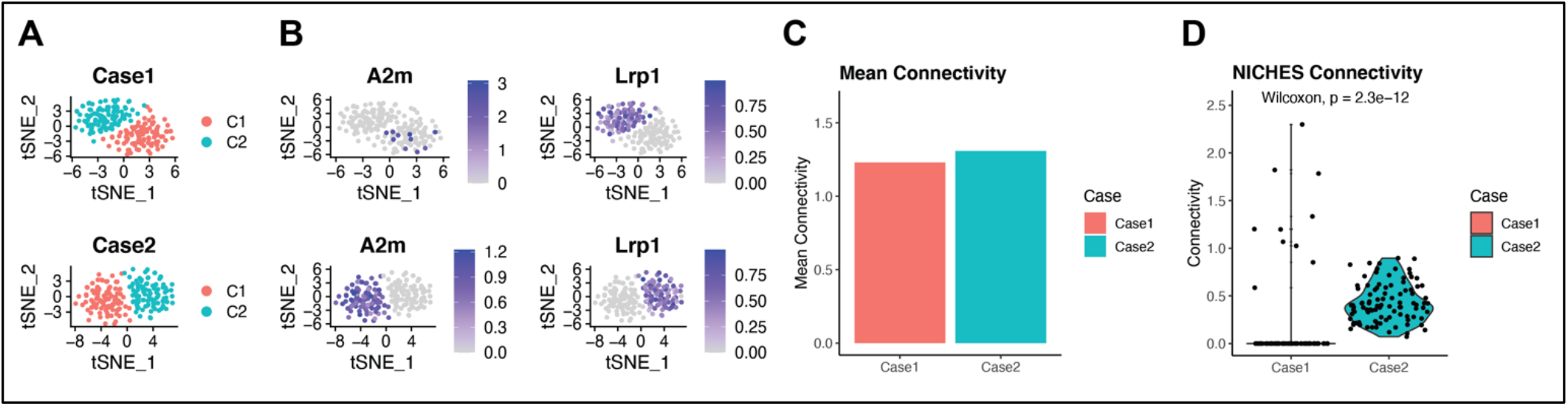
NICHES captures cross-condition archetype shift in cell-cell connectivity. In this simulation using synthetic data (see Methods), we compare two cell-systems (A, Case 1 and Case 2) representing different experimental conditions or tissues containing the same number of cells, with the same celltypes present, and similar mean connectivity for a given signaling mechanism. Case 1 cells express ligand sparsely but highly, while Case 2 sending cells express ligand broadly but lowly (B). Receptor expression is identical in each case. While mean connectivity is nearly identical (C), NICHES captures the significantly different ground-truth connectivity between the two cases (D).

**Figure S4:**
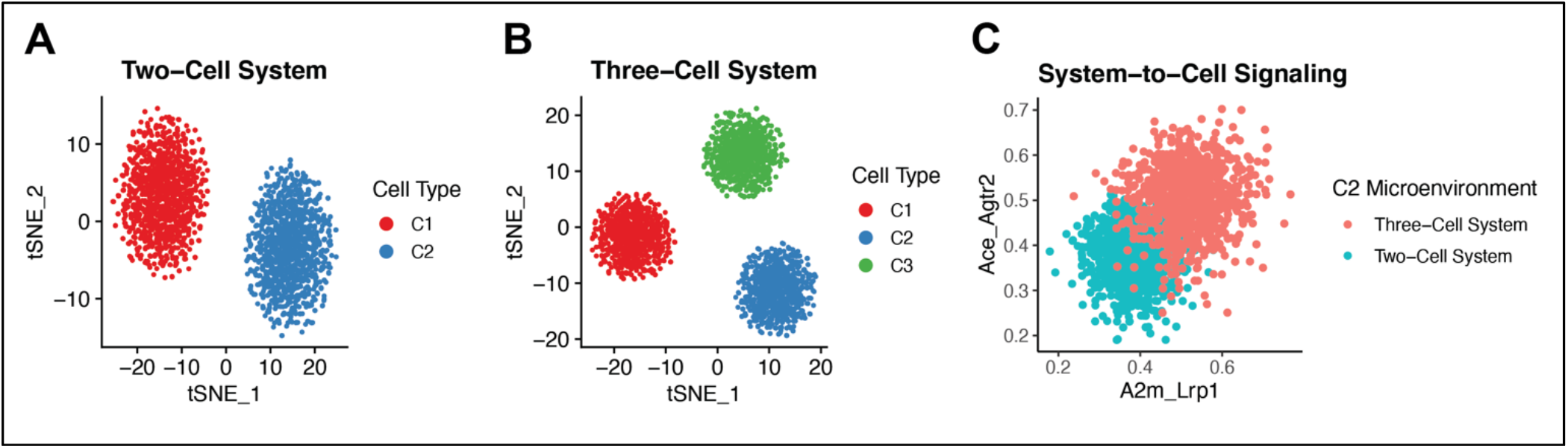
Differential System-to-Cell Signaling. In this simulation using synthetic data (see Methods), we demonstrate the capability of NICHES to measure altered system-cell signaling topology due to the addition or removal of cells. In Case 1 we have a two-cell system containing communicating cell types C1 and C2 (A). In Case 2, a third cell type (C3) has entered the system which expresses ligands cognate to receptors on C1 and C2 (B). When we use NICHES to calculate system-to-cell signaling within each case, we see a clear shift in the character of the sensed environment of celltype C2 due to the altered mean ligand expression within the system. This functionality of NICHES empowers the study of complex biological questions, such as how added, aberrant or infiltrating cells might affect the microenvironment of a receiving celltype across experimental conditions or disease states. A biological counterpart to this simulation is shown here: https://msraredon.github.io/NICHES/articles/09%20System%20Effects%20of%20Aberrant%20Cells.html

**Figure S5:**
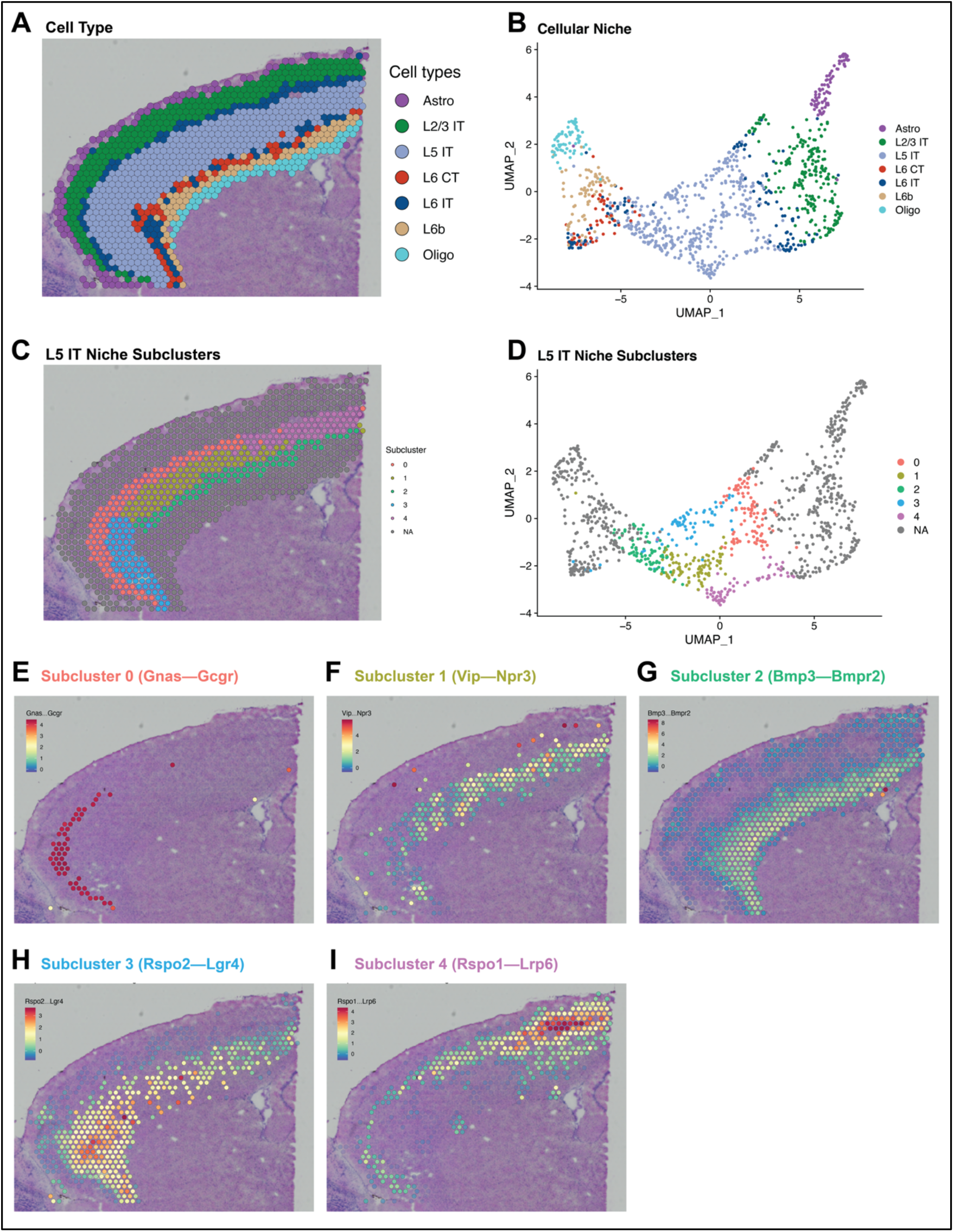
NICHES Reveals Intra-Celltype Microenvironment Heterogeneity. A) Spatial transcriptomic data labeled by dominant celltype (see Methods). B) UMAP embedding of cellular niche for each transcriptomic location. C) Sub-clustering of the L5 IT niche represented spatially and D) within UMAP space. Exploration of marker mechanisms reveals niche interactions specific to the microenvironments within each subcluster (E-I).

**Figure S6:**
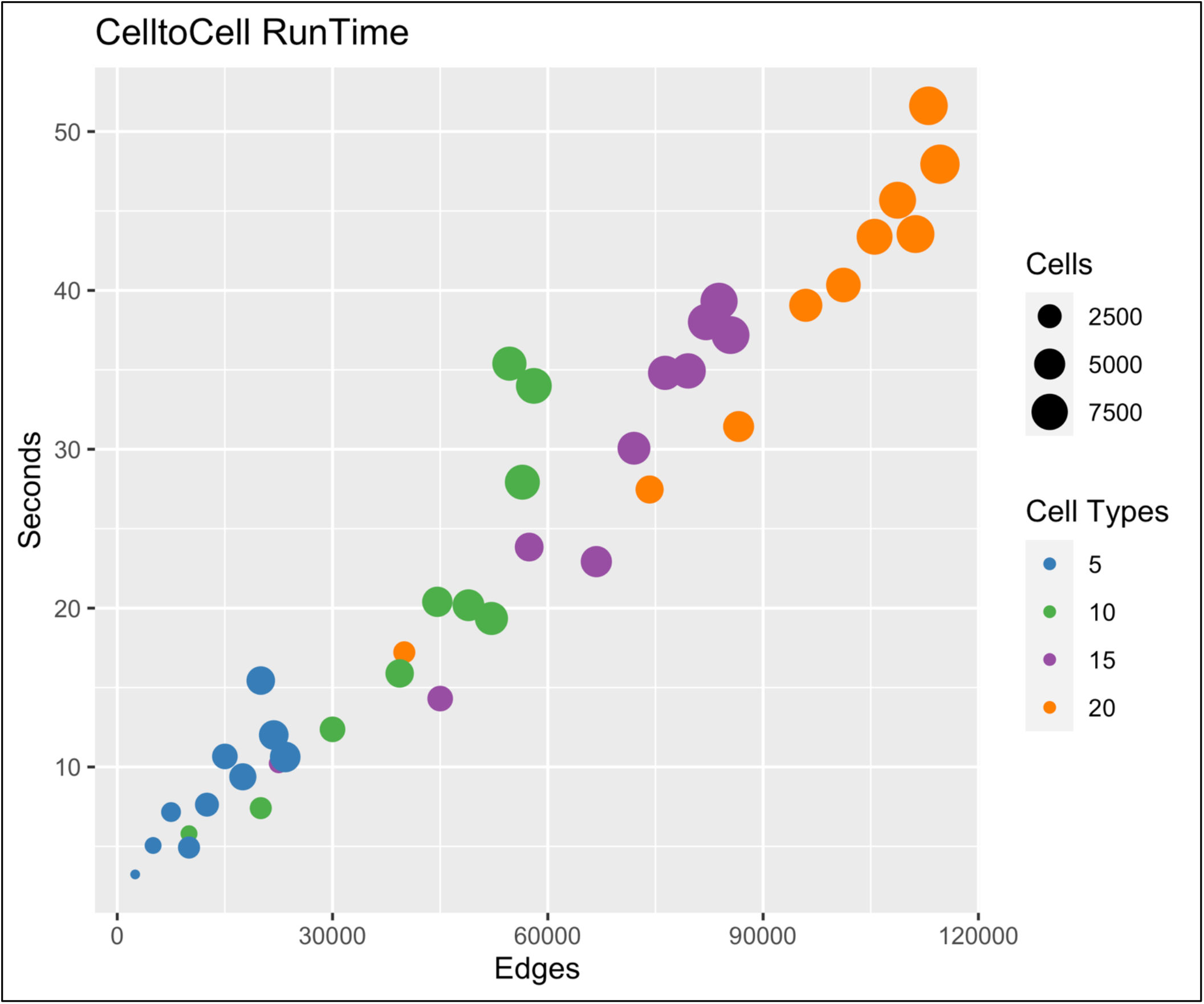
NICHES runtime and scalability. NICHES is designed to allow rapid analysis of cell-cell signaling patterns. Runtime scales reasonably well with respect to the number of edges (columns in NICHES output matrices.) Edge number is a function of input cell number and cell type number and is dataset specific.

### Software Methods

#### Dependencies

The internal workings of NICHES are dependent on a large number of other R software packages, in particular dplyr (Wickham, et al., 2019) and Seurat (Butler, et al., 2018). A maintained list of dependencies can be viewed in the DESCRIPTION file.

#### Mathematical and Computational Formalism

First, we define basic notations:

**Table S1.**
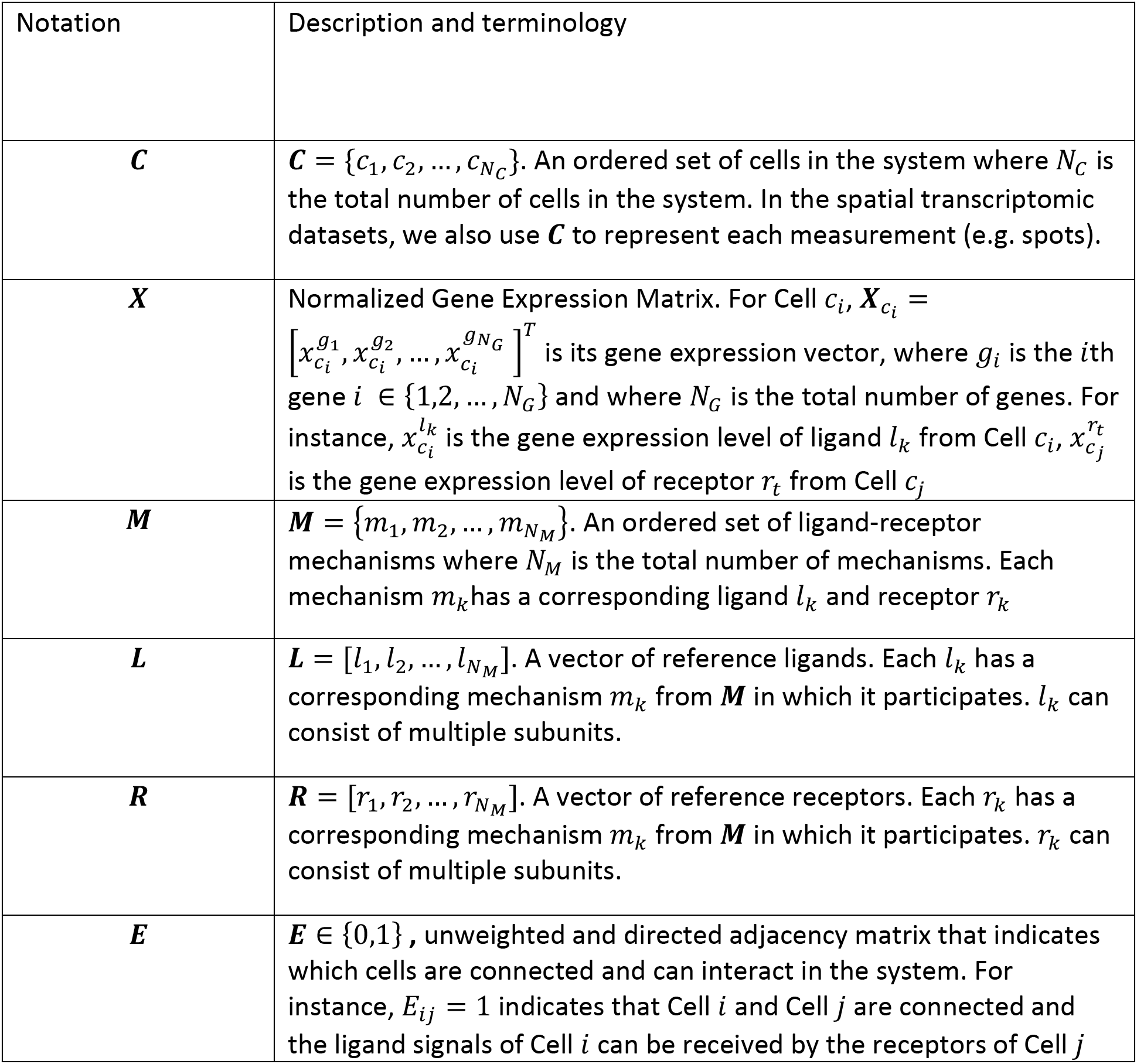
Notations

Given the gene expression data ***X*** of a cell system ***C***, along with a list of known ligand-receptor mechanism ***M***, we aim to define a vector 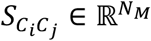 for every connected cell pair cell *i* and cell *j* in a pre-defined cell adjacency matrix *E*, such that *S_C_i_C_j__* can characterize the *N_M_* -dimensional ligand-receptor interaction profiles between cell *i* sending signal via ligand and cell *j* receiving signal via its receptors.

#### Cell-Cell Matrix Construction

To construct the Cell-Cell Matrix, we define 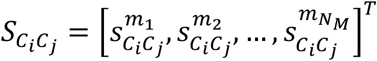 in which 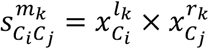, where the mechanism *m_k_* consists of ligand *l_k_* and receptor *r_k_*. We choose the multiplication operation so when the ligand or the receptor are not expressed the product is zero, representing no cell-cell signaling.

We concatenate the *S_C_i_C_j__* Cell-Cell Interaction vectors to construct the Cell-Cell Matrix: 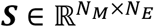, where *N_M_* is the total number of mechanisms and 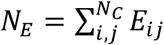 is the total number of (directed) connected cells.

***S*** can be used as input to many computational analysis pipelines, including dimensionality reduction, clustering, differential expression, pseudo-temporal ordering, and trajectory inference, etc., to investigate cell-cell interactions at the individual cell level.

#### Computing the Adjacency Matrix **E**

One step before computing a Cell-Cell Matrix is to compute the adjacency matrix ***E***. For single-cell RNA-seq datasets we assume a fully connected cellular system. However, the computational complexity of ***S*** becomes 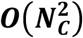, which greatly hinders the application of Cell-Cell Matrix onto cellular systems of large number of cells (e.g. *N_C_* > 1 × 10^3^).

To reduce the complexity, we adopt a random sampling scheme to down-sample edges and to compute a new 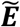 as follows: Let’s denote the set of cell type labels in the system by *P* = {*p*_1_, *p*_2_, …, *p_N_P__*} where *N_P_* is the total number of cell types. The set of cells associated with each type is denoted by *N* = {*n*_1_, *n*_2_, …, *n_N_p__*}. For each pair of cell types within {(*p_k_, p_m_*)|*k* = 1,2, …, *N_P_*; *m* = 1,2, …, *N_P_*}, we draw 2 sets of cells *C^sub,p_k_^* and *C^sub,P_m_^* from cells of cell type *p_k_* and *p_m_* uniformly, i.e., {*C^*sub,p_k_*^* ⊆ *C^P_k_^*| |*C^*sub,p_k_*^*| = min (*n_k_, n_m_*)} and {*C^*sub, p_m_*^* ⊆ *C^p_m_^* | |*C^*sub, P_m_*^* | = min (*n_k_, n_m_*)}(|*S*| denotes the number of elements in set S). Then we pair up *C^*sub, p_k_*^* and 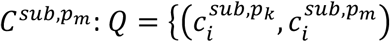|*i* = 1,2, …, min (*n_k_, n_m_*)}. Lastly, each entry in the new adjacency matrix can be set as 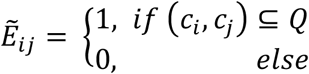.

For spatial transcriptomic datasets we constrain the interactions among cells to be only within a certain distance threshold or a certain local neighborhood. To be more specific, let us denote ***D*** as the Euclidean distance matrix among cells computed from the spatial locations, where *d_ij_* is the distance between Cell *i* and Cell *j*. Given a distance threshold *r, E_ij_* is computed as 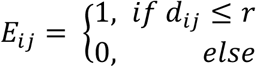 for each entry of ***E*** for spatial transcriptomic datasets. Alternatively, the user can specify the parameter k which computes a k-nearest neighbor (knn) graph from ***D*** and the adjacency matrix *E* will be computed as a mutual nearest neighbor graph from this knn graph, i.e., as 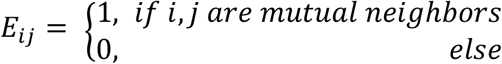

#### Niche Matrix Construction

Besides the base cell-cell interaction formulation, we extend our original definition of the Cell-Cell Matrix to investigate cellular niche and cellular influence interactions.

Specifically, we define the Niche Matrix as: 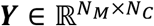, where *N_M_* is the total number of mechanisms and *N_C_* is the total number of cells in the system. A column vector of ***Y*** is defined as 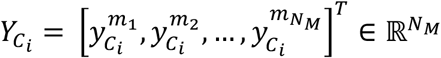, i.e., each column of ***Y*** is a *N_M_* -dimensional vector that characterizes the interaction profiles between cells sending ligand signals to Cell *i* which possesses the relevant receptors to receive these signals.

The connectivity value on one mechanism (e.g. *m_k_*) between sending cells and Cell *i* is defined as 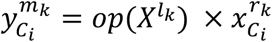 where the mechanism *m_k_* consists of ligand *l_k_* and receptor *r_k_*, *X^l_k_^* denotes the row vector of *l_k_*’s expression levels across the cells that are connected to Cell *i*, and *op*() is a vector operator which, in our implementation, can be either *sum* (default) or *mean*.

Similarly, we define the Influence Matrix as: 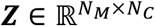 where each column of ***Z*** is a *N_M_* -dimensional vector that characterizes the interaction profiles between Cell *i* that sends the ligand signals and the cells receiving from it. Each connectivity value between Cell *i* and the system is defined as 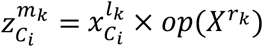 where the mechanism *m_k_* consists of ligand *l_k_* and receptor *r_k_*, *X^r^k* denotes the row vector of *r_k_*’s expression levels across the cells that connect to Cell *i* in the system, and *op*() is again either *sum* (default) or *mean*.

For single-cell RNA-seq datasets without spatial coordinates, we assume a fully connected ***E*** involving all cells measured within a system. For spatial transcriptomic datasets, we construct ***E*** in the same fashion as for the spatial Cell-Cell Matrix, limiting edges to neighbors within radius *r* or within a user-defined set of nearest neighbors.

#### Metadata Mapping

NICHES allows the researcher to carry over any and all metadata (i.e. sample labels, coarse- and fine-grain cluster labeling, experimental conditions) from source data, allowing rapid downstream differential analysis between already tagged groupings of cells. For every input metadata category, the NICHES Cell-Cell Matrix output object has Sending Metadata, Receiving Metadata, and Sending-Receiving Metadata associated with every column. The Niche Matrix and Influence Matrix have only Receiving Metadata and Sending Metadata associated with their columns, respectively.

Because each Cell-Cell Matrix contains many individual measurements of cell pairings (or environment-cell pairings in the Niche Matrix), differential analysis can be used to reveal ligand-receptor mechanisms preferential to a given celltype-celltype cross within a system, or to identify top differential signaling mechanisms across subject, disease state, experimental condition, or tissue. Such calculations may be performed for specific celltype-celltype crosses or for other custom groupings as the user desires, based on mapped metadata. We recommend using ROC analysis to measure how well a mechanism differentiates two groups compared to standard two-sample tests when the columns in Cell-Cell Matrix or Niche Matrix are no longer independent.

### Application Methods

#### Methods for Application to Synthetic Data

We generate 3 simulation datasets (*Simulation 1, Simulation 2, Simulation 3*) for 3 separate simulation analyses (Supplemental Fig A-C, Fig E-I, Fig J-L) respectively. For each dataset, we simulate 3 categories of genes: signaling genes (ligands and receptors), 50 non-signaling marker genes to differentiate each cell type, and 5000 noise genes. We assume the genes in the datasets follow negative binomial (NB) distributions parametrized by parameter *μ* which characterizes the mean expression level, and the dispersion parameter *γ*. We describe the exact parameter settings for each dataset as follows:

**Table S2:**
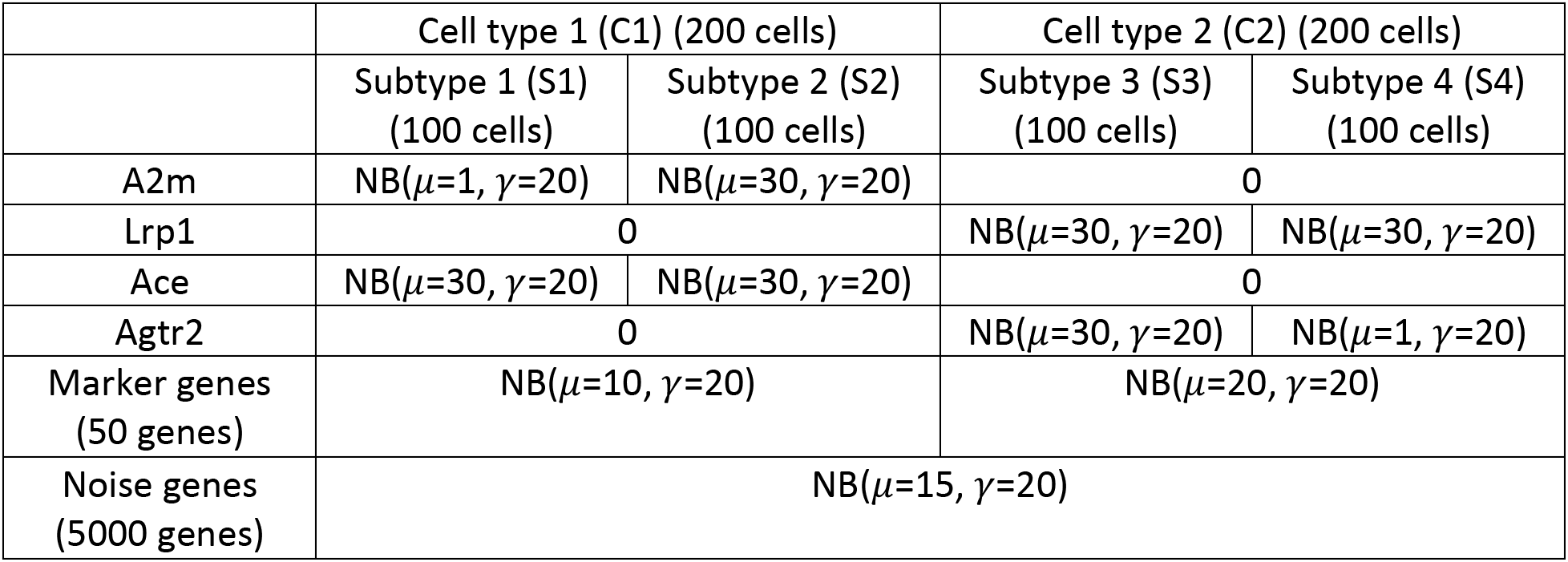
Count matrix design for *Simulation 1*

**Table S3:**
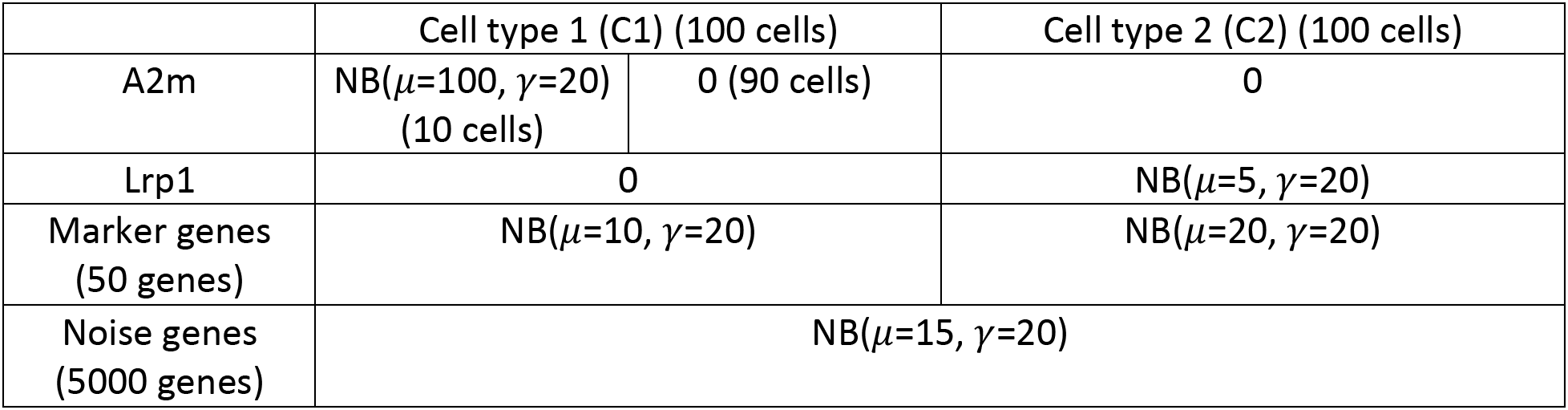
Count matrix design for *Simulation 2* (*Case 1*)

**Table S4:**
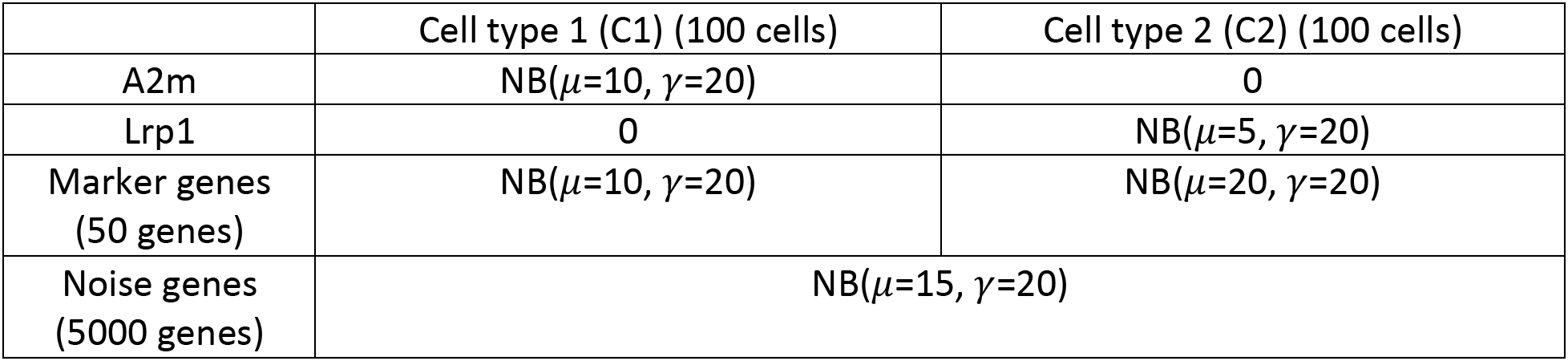
Count matrix design for *Simulation 2* (*Case 2*)

**Table S5:**
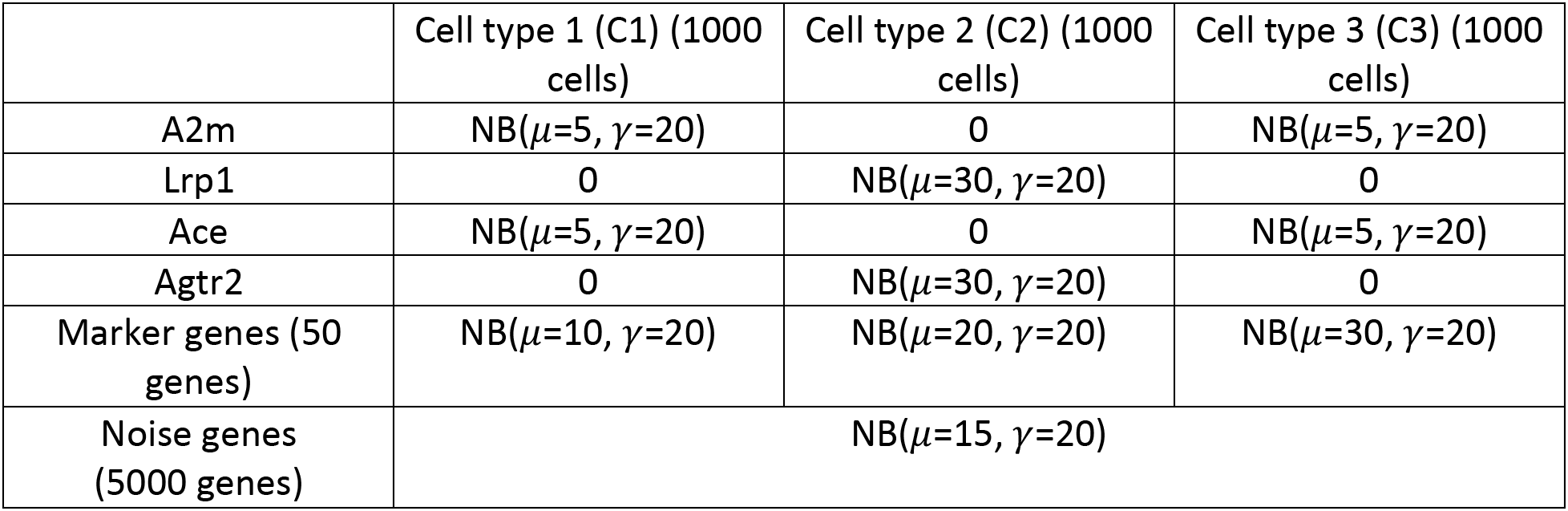
Count matrix design for *Simulation 3*

#### Methods for Application to Native Lung scRNAseq Data

Data was downloaded from (Raredon, et al., 2019), subset to 4 main populations of interest, and run through standard principle component analysis (PCA), clustering, and UMAP embedding pipelines in Seurat (McInnes, et al., 2018; Stuart, et al., 2019). Data was imputed using ALRA (Linderman, et al., 2022) and then run through the NICHES function RunCellToCell. The resulting signaling matrix was then used to create a new Seurat object which was scaled and run through PCA again and embedding using UMAP. FindAllMarkers was used in Seurat to identify cell-cell interaction markers of interest.

#### Methods for Application to Brain Spatial Transcriptomic Data

Anterior mouse brain data was downloaded from 10× Genomics (2020) and preprocessed following the steps in Seurat (Stuart, et al., 2019), with subsetting to the frontal cortex region only. We integrated the data with a reference single-cell RNA-seq dataset (Tasic, et al., 2016) and used its cell type annotations to predict the labels of the spatial pixels by a probabilistic classifier (Seurat TransferData function). We then annotated each spatial pixel by its most probable label.

For NICHES matrix construction, we imputed the data with ALRA (Linderman, et al., 2022) based on the normalized data matrix, and then applied the NICHES Neighborhood-to-Cell function to compute niche signaling between direct histologic neighbors. The resulting niche matrix was embedded using UMAP (McInnes, et al., 2018) in Seurat 4.0 (Hao, et al., 2021). FindAllMarkers in Seurat was used to compute top markers for each celltype niche.

