## Supplementary Materials for "Comprehensive visualization of cell-cell interactions in single-cell and spatial transcriptomics with NICHES"

**This PDF files includes:**

Supplemental Text

Supplemental Findings

Supplemental Figures

Software Methods

Application Methods

Supplemental References

**Software and Vignettes associated with this manuscript are available here:**

https://msraredon.github.io/NICHES/

https://github.com/msraredon/NICHES

10.5281/zenodo.6846813

**Data used in Vignettes is available at:**

10.5281/zenodo.6846617

10.5281/zenodo.6878944

10.5281/zenodo.6878953

**Supplemental Text**

### Supplemental Figures

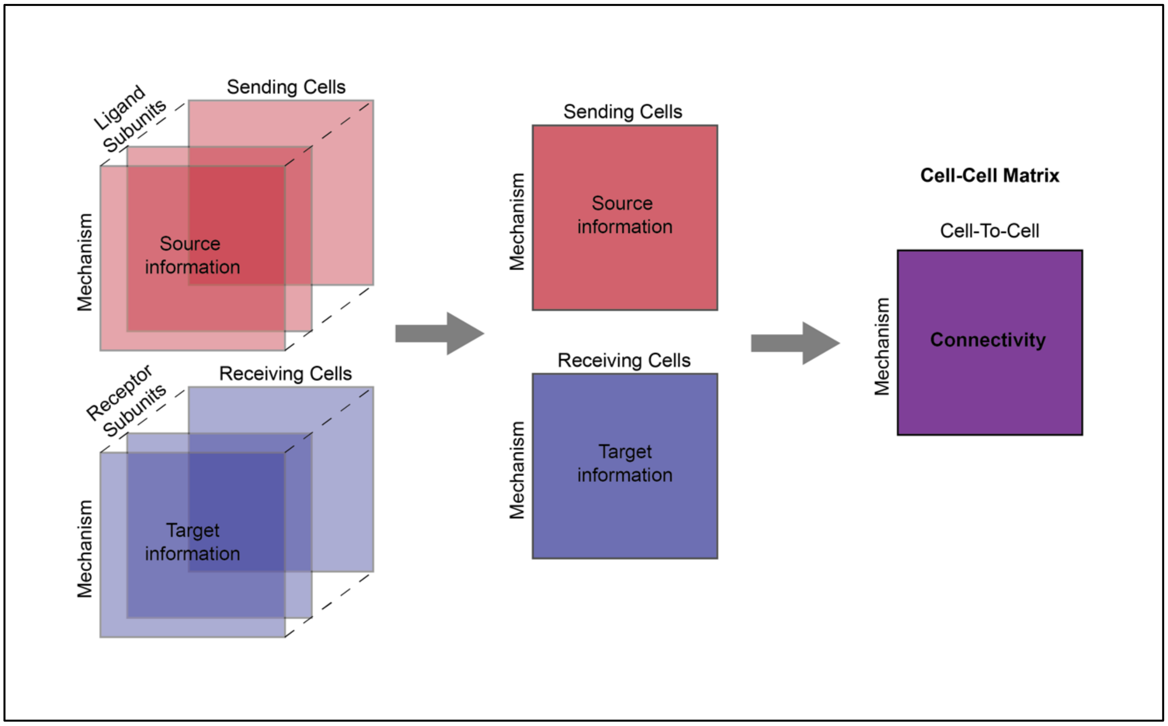


**Figure S1: NICHES connectivity for mechanisms with multiple subunits**

NICHES is capable of calculating intercellular connectivity for ligand-receptor mechanisms with any number of subunits. For a given mechanism, ligand subunit expressivities on the sending cell are multiplied together and receptor subunit expressivities on the receiving cell are multiplied together. These two values are then multiplied to yield mechanism connectivity between the sending and receiving cell. This operation is zero-preserving by design, so that the lack of expression of even a single subunit on either the sending or receiving cell within a given cell-cell pairing will cause a connectivity value of zero.

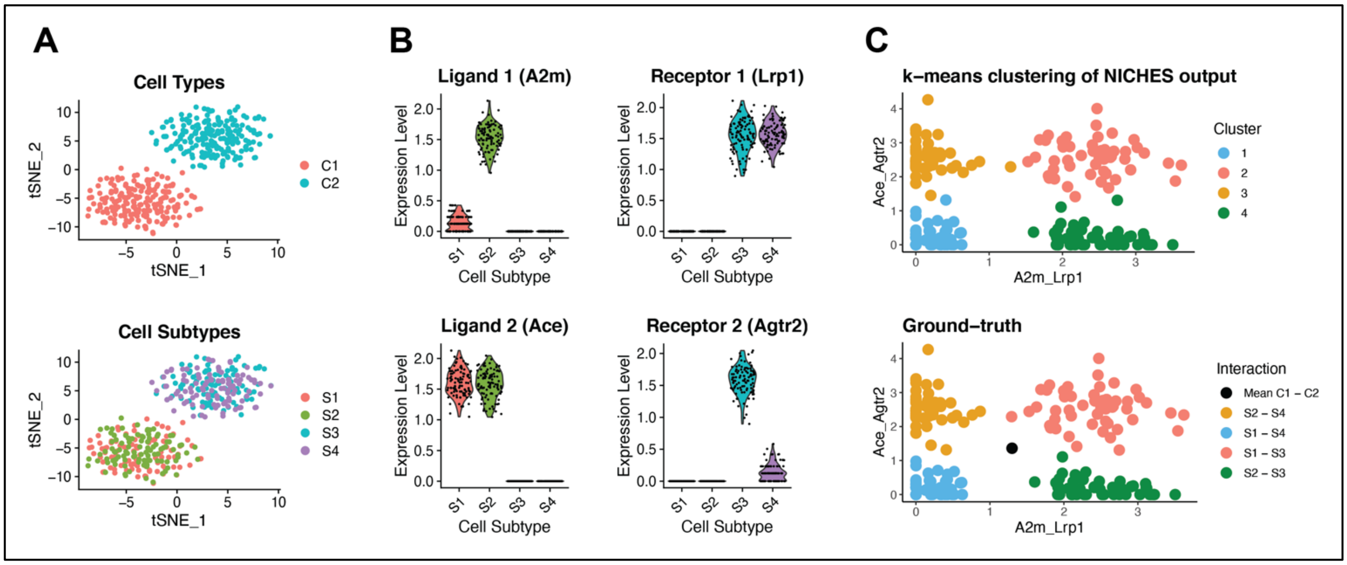


**Figure S2: NICHES captures heterogeneity in cell-cell connectivity**

In this simulation using synthetic data (see Methods) there are two celltypes labeled C1 and C2 containing signaling subtypes S1-S4 which do not resolve in gene space (A). These subtypes communicate in distinct ways via two distinct signaling mechanisms: A2m-Lrp1 and Ace-Agtr2 (B). Subpopulation S2 expresses ligand A2m higher than S1 while subpopulation S3 expresses receptor Agtr2 higher than S4. This expression pattern creates four distinct cell-cell signaling relationships even though only two celltypes have been crossed. NICHES allows rapid observation of these distinct relationships using two-dimensional embeddings and k-means clustering (C, top) which closely matches the ground truth subtype crosses in this simulation (C, bottom). Mean connectivity between C1 and C2 is represented in black in the lower panel of (C). A biological counterpart to this simulation is show in Figure 1F.

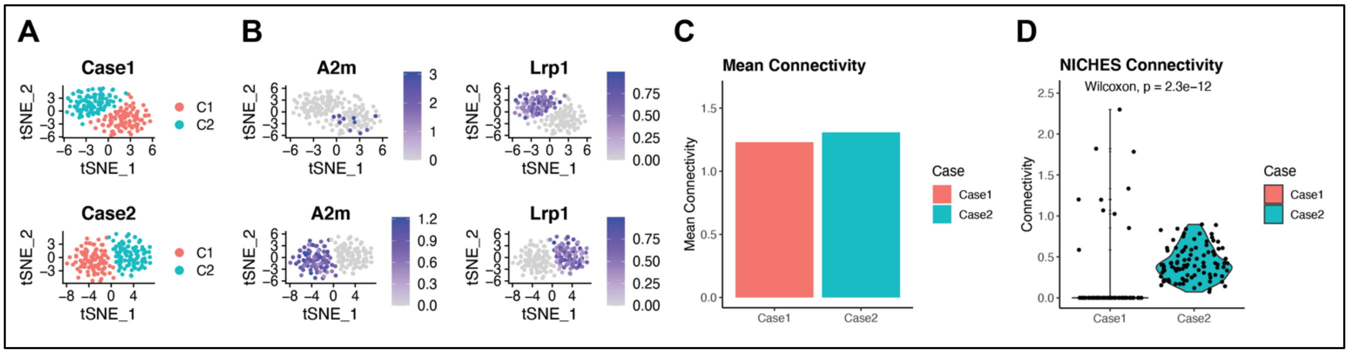


**Figure S3: NICHES captures cross-condition archetype shift in cell-cell connectivity**

In this simulation using synthetic data (see Methods), we compare two cell-systems (A, Case 1 and Case 2) representing different experimental conditions or tissues containing the same number of cells, with the same celltypes present, and similar mean connectivity for a given signaling mechanism. Case 1 cells express ligand sparsely but highly, while Case 2 sending cells express ligand broadly but lowly (B). Receptor expression is identical in each case. While mean connectivity is nearly identical (C), NICHES captures the significantly different ground-truth connectivity between the two cases (D).

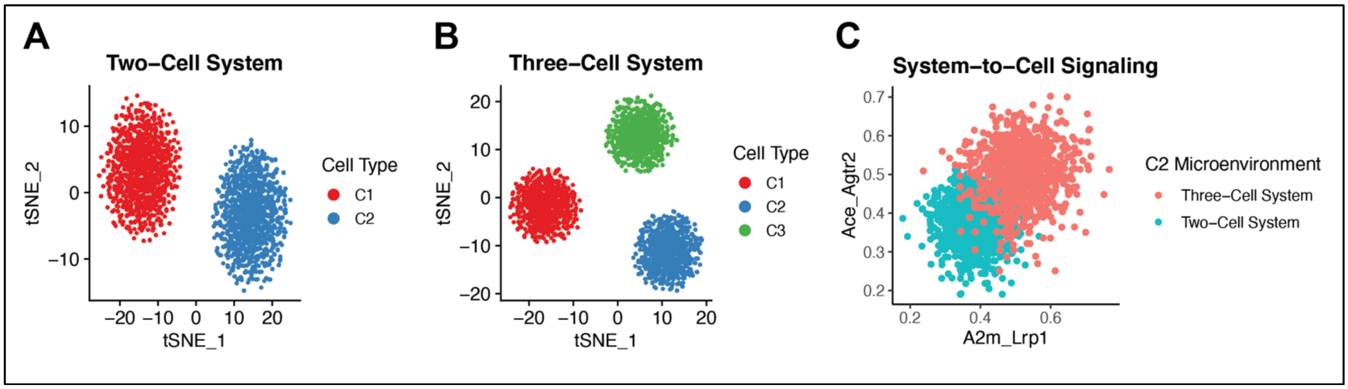


**Figure S4: Differential System-to-Cell Signaling**

In this simulation using synthetic data (see Methods), we demonstrate the capability of NICHES to measure altered system-cell signaling topology due to the addition or removal of cells. In Case 1 we have a two-cell system containing communicating cell types C1 and C2 (A). In Case 2, a third cell type (C3) has entered the system which expresses ligands cognate to receptors on C1 and C2 (B). When we use NICHES to calculate system-to-cell signaling within each case, we see a clear shift in the character of the sensed environment of celltype C2 due to the altered mean ligand expression within the system. This functionality of NICHES empowers the study of complex biological questions, such as how added, aberrant or infiltrating cells might affect the microenvironment of a receiving celltype across experimental conditions or disease states. A biological counterpart to this simulation is shown here: https://msraredon.github.io/NICHES/articles/09%20System%20Effects%20of%20Aberrant%20Cells.html

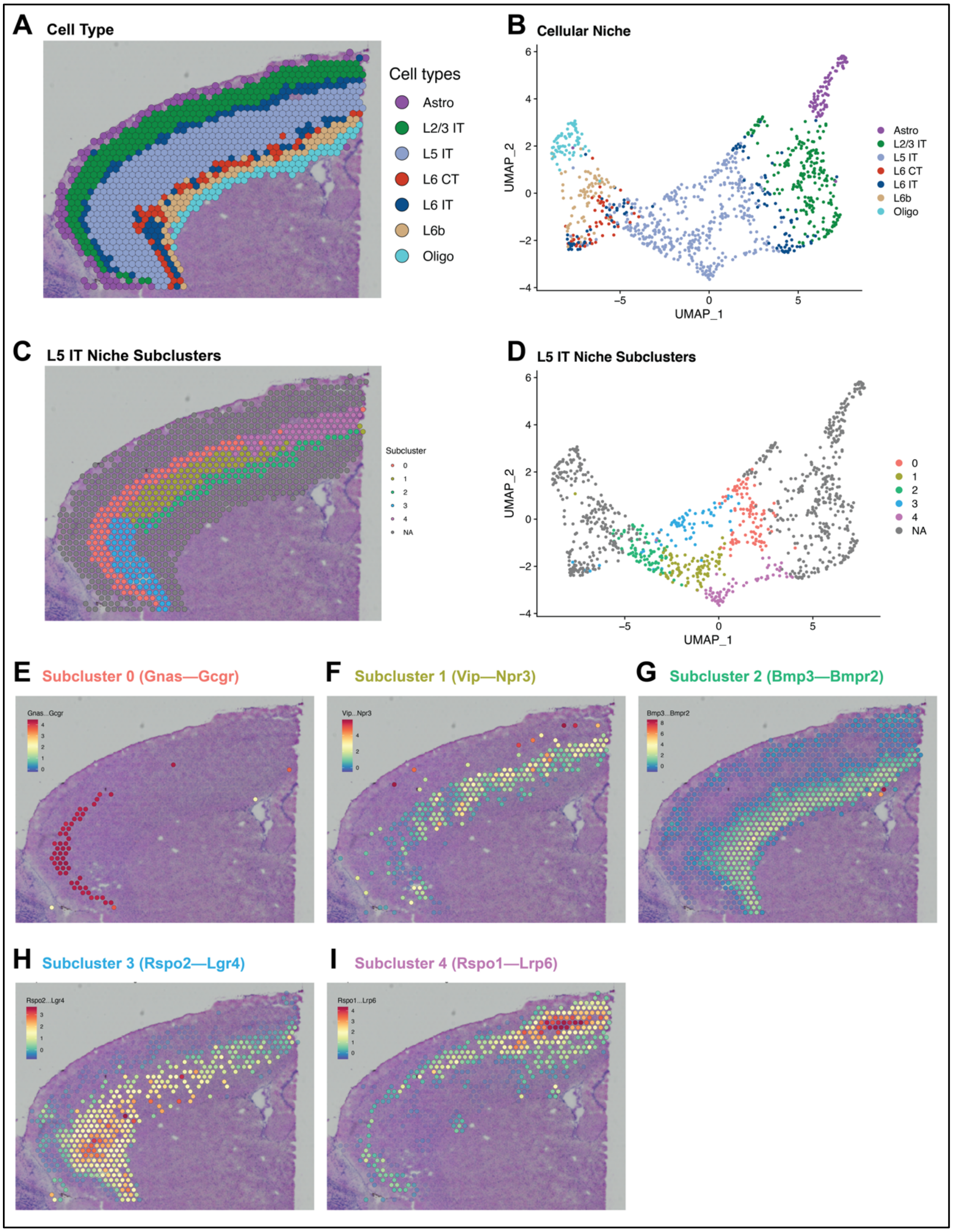


**Figure S5: NICHES Reveals Intra-Celltype Microenvironment Heterogeneity**

A) Spatial transcriptomic data labeled by dominant celltype (see Methods). B) UMAP embedding of cellular niche for each transcriptomic location. C) Sub-clustering of the L5 IT niche represented spatially and D) within UMAP space. Exploration of marker mechanisms reveals niche interactions specific to the microenvironments within each subcluster (E-I).

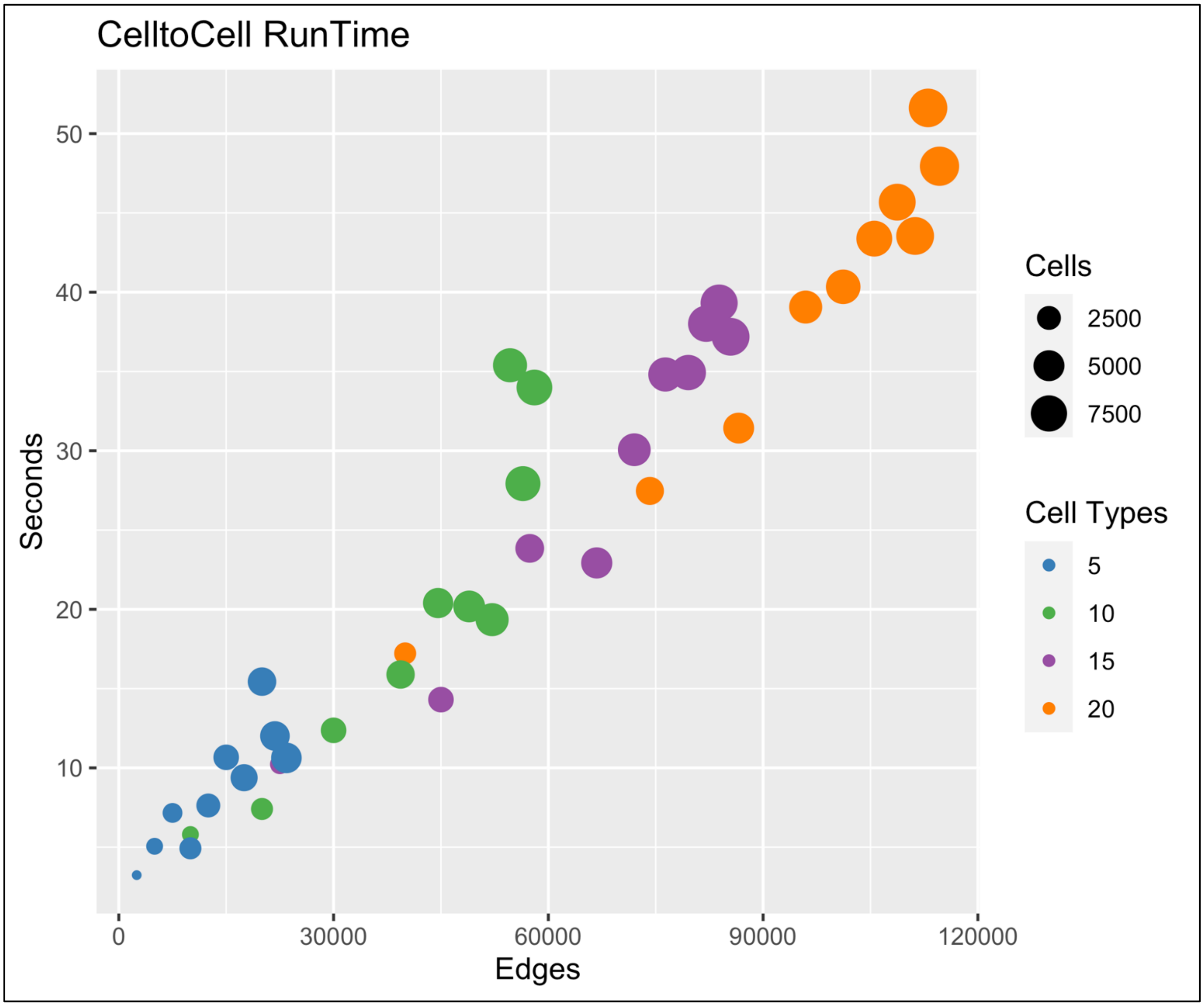


**Figure S6: NICHES runtime and scalability**

NICHES is designed to allow rapid analysis of cell-cell signaling patterns. Runtime scales reasonably well with respect to the number of edges (columns in NICHES output matrices.) Edge number is a function of input cell number and cell type number and is dataset specific.

**Software Methods**

Dependencies

The internal workings of NICHES are dependent on a large number of other R software packages, in particular dplyr (Wickham, et al., 2019) and Seurat (Butler, et al., 2018). A maintained list of dependencies can be viewed in the DESCRIPTION file.

Mathematical and Computational Formalism

First, we define basic notations:

Table S1. Notations

| Notation | Description and terminology |
|---|---|
| $\boldsymbol{C}$ | $\boldsymbol{C} = \{c_1, c_2, \dots, c_{N_C}\}$. An ordered set of cells in the system where $N_C$ is the total number of cells in the system. In the spatial transcriptomic datasets, we also use $\boldsymbol{C}$ to represent each measurement (e.g. spots). |
| $\boldsymbol{X}$ | Normalized Gene Expression Matrix. For Cell $c_i$, $\boldsymbol{X}_{c_i} = \left[x_{c_i}^{g_1}, x_{c_i}^{g_2}, \dots, x_{c_i}^{g_{N_G}}\right]^T$ is its gene expression vector, where $g_i$ is the $i$th gene $i \in \{1,2, \dots, N_G\}$ and where $N_G$ is the total number of genes. For instance, $x_{c_i}^{l_k}$ is the gene expression level of ligand $l_k$ from Cell $c_i$, $x_{c_j}^{r_t}$ is the gene expression level of receptor $r_t$ from Cell $c_j$ |
| $\boldsymbol{M}$ | $\boldsymbol{M} = \{m_1, m_2, \dots, m_{N_M}\}$. An ordered set of ligand-receptor mechanisms where $N_M$ is the total number of mechanisms. Each mechanism $m_k$has a corresponding ligand $l_k$ and receptor $r_k$ |
| $\boldsymbol{L}$ | $\boldsymbol{L} = [l_1, l_2, \dots, l_{N_M}]$. A vector of reference ligands. Each $l_k$ has a corresponding mechanism $m_k$ from $\boldsymbol{M}$ in which it participates. $l_k$ can consist of multiple subunits. |
| $\boldsymbol{R}$ | $\boldsymbol{R} = [r_1, r_2, \dots, r_{N_M}]$. A vector of reference receptors. Each $r_k$ has a corresponding mechanism $m_k$ from $\boldsymbol{M}$ in which it participates. $r_k$ can consist of multiple subunits. |
| $\boldsymbol{E}$ | $\boldsymbol{E} \in \{0,1\}$ **,** unweighted and directed adjacency matrix that indicates which cells are connected and can interact in the system. For instance, $E_{ij} = 1$ indicates that Cell $i$ and Cell $j$ are connected and the ligand signals of Cell $i$ can be received by the receptors of Cell $j$ |

Given the gene expression data $\boldsymbol{X}$ of a cell system $\boldsymbol{C}$, along with a list of known ligand-receptor mechanism $\boldsymbol{M}$, we aim to define a vector $S_{C_iC_j} \in \mathbb{R}^{N_M}$ for every connected cell pair cell $i$ and cell $j$ in a pre-defined cell adjacency matrix $\boldsymbol{E}$, such that $S_{C_iC_j}$ can characterize the $N_M$ -dimensional ligand-receptor interaction profiles between cell $i$ sending signal via ligand and cell $j$ receiving signal via its receptors.

Cell-Cell Matrix Construction

To construct the Cell-Cell Matrix, we define $S_{C_iC_j} = \left[s_{C_iC_j}^{m_1}, s_{C_iC_j}^{m_2}, \dots, s_{C_iC_j}^{m_{N_M}}\right]^T$ in which $s_{C_iC_j}^{m_k} = x_{C_i}^{l_k} \times x_{C_j}^{r_k}$, where the mechanism $m_k$ consists of ligand $l_k$ and receptor $r_k$. We choose the multiplication operation so when the ligand or the receptor are not expressed the product is zero, representing no cell-cell signaling.

We concatenate the $S_{C_iC_j}$ Cell-Cell Interaction vectors to construct the Cell-Cell Matrix: $\boldsymbol{S} \in \mathbb{R}^{N_M \times N_E}$, where $N_M$ is the total number of mechanisms and $N_E = \sum_{i,j}^{N_C} E_{ij}$ is the total number of (directed) connected cells.

$\boldsymbol{S}$ can be used as input to many computational analysis pipelines, including dimensionality reduction, clustering, differential expression, pseudo-temporal ordering, and trajectory inference, etc., to investigate cell-cell interactions at the individual cell level.

Computing the Adjacency Matrix $\mathbf{E}$

One step before computing a Cell-Cell Matrix is to compute the adjacency matrix $\boldsymbol{E}$. For single-cell RNA-seq datasets we assume a fully connected cellular system. However, the computational complexity of $\boldsymbol{S}$ becomes $\boldsymbol{O}\left(\boldsymbol{N_C^2}\right)$**,** which greatly hinders the application of Cell-Cell Matrix onto cellular systems of large number of cells (e.g. $N_C > 1 \times 10^3$).

To reduce the complexity, we adopt a random sampling scheme to down-sample edges and to compute a new $\widetilde{\boldsymbol{E}}$ as follows: Let's denote the set of cell type labels in the system by $P = \{p_1, p_2, \dots, p_{N_P}\}$ where $N_P$ is the total number of cell types. The set of cells associated with each type is denoted by $N = \{n_1, n_2, \dots, n_{N_P}\}$. For each pair of cell types within $\{(p_k, p_m) | k = 1,2, \dots, N_P; m = 1,2, \dots, N_P\}$, we draw 2 sets of cells $C^{sub,p_k}$and $C^{sub,p_m}$ from cells of cell type $p_k$ and $p_m$ uniformly, i.e., $\{C^{sub,p_k} \subseteq C^{p_k} |\, |C^{sub,p_k}| = \min{(n_k, n_m)}\}$ and $\{C^{sub,p_m} \subseteq C^{p_m} |\, |C^{sub,p_m}| = \min{(n_k, n_m)}\}$ ($|S|$ denotes the number of elements in set S). Then we pair up $C^{sub,p_k}$ and $C^{sub,p_m}$: $Q = \left\{\left(c_i^{sub,p_k}, c_i^{sub,p_m}\right) \middle| i = 1,2, \dots, \min{(n_k, n_m)}\right\}$. Lastly, each entry in the new adjacency matrix can be set as $\tilde{E}_{ij} = \begin{cases} 1, & if\ (c_i, c_j) \subseteq Q \\ 0, & else \end{cases}$.

For spatial transcriptomic datasets we constrain the interactions among cells to be only within a certain distance threshold or a certain local neighborhood. To be more specific, let us denote $\boldsymbol{D}$ as the Euclidean distance matrix among cells computed from the spatial locations, where $d_{ij}$ is the distance between Cell $i$ and Cell $j$. Given a distance threshold $r$, $E_{ij}$ is computed as $E_{ij} = \begin{cases} 1, & if\ d_{ij} \leq r \\ 0, & else \end{cases}$ for each entry of $\boldsymbol{E}$ for spatial transcriptomic datasets. Alternatively, the user can specify the parameter k which computes a k-nearest neighbor (knn) graph from $\boldsymbol{D}$ and the adjacency matrix $E$ will be computed as a mutual nearest neighbor graph from this knn graph, i.e., as $E_{ij} = \begin{cases} 1, & if\ i, j\ are\ mutual\ neighbors \\ 0, & else \end{cases}$

Niche Matrix Construction

Besides the base cell-cell interaction formulation, we extend our original definition of the Cell-Cell Matrix to investigate cellular niche and cellular influence interactions.

Specifically, we define the Niche Matrix as: $\boldsymbol{Y} \in \mathbb{R}^{N_M \times N_C}$, where $N_M$ is the total number of mechanisms and $N_C$ is the total number of cells in the system. A column vector of $\boldsymbol{Y}$ is defined as $Y_{C_i} = \left[y_{C_i}^{m_1}, y_{C_i}^{m_2}, \ldots, y_{C_i}^{m_{N_M}}\right]^T \in \mathbb{R}^{N_M}$, i.e., each column of $\boldsymbol{Y}$ is a $N_M$ -dimensional vector that characterizes the interaction profiles between cells sending ligand signals to Cell $i$ which possesses the relevant receptors to receive these signals.

The connectivity value on one mechanism (e.g. $m_k$) between sending cells and Cell $i$ is defined as $y_{C_i}^{m_k} = op(X^{l_k}) \times x_{C_i}^{r_k}$ where the mechanism $m_k$ consists of ligand $l_k$ and receptor $r_k$, $X^{l_k}$ denotes the row vector of $l_k$'s expression levels across the cells that are connected to Cell $i$, and $op()$ is a vector operator which, in our implementation, can be either $sum$ (default) or $mean$.

Similarly, we define the Influence Matrix as: $\boldsymbol{Z} \in \mathbb{R}^{N_M \times N_C}$ where each column of $\boldsymbol{Z}$ is a $N_M$ -dimensional vector that characterizes the interaction profiles between Cell $i$ that sends the ligand signals and the cells receiving from it. Each connectivity value between Cell $i$ and the system is defined as $z_{C_i}^{m_k} = x_{C_i}^{l_k} \times op(X^{r_k})$ where the mechanism $m_k$ consists of ligand $l_k$ and receptor $r_k$, $X^{r_k}$ denotes the row vector of $r_k$'s expression levels across the cells that connect to Cell $i$ in the system, and $op()$ is again either $sum$ (default) or $mean$.

For single-cell RNA-seq datasets without spatial coordinates, we assume a fully connected $\boldsymbol{E}$ involving all cells measured within a system. For spatial transcriptomic datasets, we construct $\boldsymbol{E}$ in the same fashion as for the spatial Cell-Cell Matrix, limiting edges to neighbors within radius $r$ or within a user-defined set of nearest neighbors.

Table S2: Count matrix design for *Simulation 1*

| | Cell type 1 (C1) (200 cells) | | Cell type 2 (C2) (200 cells) | |
|---|---|---|---|---|
| | Subtype 1 (S1) (100 cells) | Subtype 2 (S2) (100 cells) | Subtype 3 (S3) (100 cells) | Subtype 4 (S4) (100 cells) |
| A2m | NB($\mu$=1, $\gamma$=20) | NB($\mu$=30, $\gamma$=20) | 0 | |
| Lrp1 | 0 | | NB($\mu$=30, $\gamma$=20) | NB($\mu$=30, $\gamma$=20) |
| Ace | NB($\mu$=30, $\gamma$=20) | NB($\mu$=30, $\gamma$=20) | 0 | |
| Agtr2 | 0 | | NB($\mu$=30, $\gamma$=20) | NB($\mu$=1, $\gamma$=20) |
| Marker genes (50 genes) | NB($\mu$=10, $\gamma$=20) | | NB($\mu$=20, $\gamma$=20) | |
| Noise genes (5000 genes) | NB($\mu$=15, $\gamma$=20) | | | |

Table S3: Count matrix design for *Simulation* 2 (Case 1)

| | Cell type 1 (C1) (100 cells) | | Cell type 2 (C2) (100 cells) |
|---|---|---|---|
| A2m | NB($\mu$=100, $\gamma$=20) (10 cells) | 0 (90 cells) | 0 |
| Lrp1 | 0 | | NB($\mu$=5, $\gamma$=20) |
| Marker genes (50 genes) | NB($\mu$=10, $\gamma$=20) | | NB($\mu$=20, $\gamma$=20) |
| Noise genes (5000 genes) | NB($\mu$=15, $\gamma$=20) | | |

Table S4: Count matrix design for *Simulation* 2 (Case 2)

| | Cell type 1 (C1) (100 cells) | Cell type 2 (C2) (100 cells) |
|---|---|---|
| A2m | NB($\mu$=10, $\gamma$=20) | 0 |
| Lrp1 | 0 | NB($\mu$=5, $\gamma$=20) |
| Marker genes (50 genes) | NB($\mu$=10, $\gamma$=20) | NB($\mu$=20, $\gamma$=20) |
| Noise genes (5000 genes) | NB($\mu$=15, $\gamma$=20) | |

Table S5: Count matrix design for *Simulation* 3

| | Cell type 1 (C1) (1000 cells) | Cell type 2 (C2) (1000 cells) | Cell type 3 (C3) (1000 cells) |
|---|---|---|---|
| A2m | NB($\mu$=5, $\gamma$=20) | 0 | NB($\mu$=5, $\gamma$=20) |
| Lrp1 | 0 | NB($\mu$=30, $\gamma$=20) | 0 |
| Ace | NB($\mu$=5, $\gamma$=20) | 0 | NB($\mu$=5, $\gamma$=20) |
| Agtr2 | 0 | NB($\mu$=30, $\gamma$=20) | 0 |
| Marker genes (50 genes) | NB($\mu$=10, $\gamma$=20) | NB($\mu$=20, $\gamma$=20) | NB($\mu$=30, $\gamma$=20) |
| Noise genes (5000 genes) | NB($\mu$=15, $\gamma$=20) | | |

Methods for Application to Native Lung scRNAseq Data

Data was downloaded from (Raredon, et al., 2019), subset to 4 main populations of interest, and run through standard principle component analysis (PCA), clustering, and UMAP embedding pipelines in Seurat (McInnes, et al., 2018; Stuart, et al., 2019). Data was imputed using ALRA (Linderman, et al., 2022) and then run through the NICHES function RunCellToCell. The resulting signaling matrix was then used to create a new Seurat object which was scaled and run through PCA again and embedding using UMAP. FindAllMarkers was used in Seurat to identify cell-cell interaction markers of interest.

Methods for Application to Brain Spatial Transcriptomic Data

Anterior mouse brain data was downloaded from 10x Genomics (2020) and preprocessed following the steps in Seurat (Stuart, et al., 2019), with subsetting to the frontal cortex region only. We integrated the data with a reference single-cell RNA-seq dataset (Tasic, et al., 2016) and used its cell type annotations to predict the labels of the spatial pixels by a probabilistic classifier (Seurat TransferData function). We then annotated each spatial pixel by its most probable label.

For NICHES matrix construction, we imputed the data with ALRA (Linderman, et al., 2022) based on the normalized data matrix, and then applied the NICHES Neighborhood-to-Cell function to compute niche signaling between direct histologic neighbors. The resulting niche matrix was embedded using UMAP (McInnes, et al., 2018) in Seurat 4.0 (Hao, et al., 2021). FindAllMarkers in Seurat was used to compute top markers for each celltype niche.
